# Limited phenological and pollinator-mediated isolation among selfing and outcrossing *Arabidopsis lyrata* populations

**DOI:** 10.1101/2019.12.17.879361

**Authors:** Courtney E. Gorman, Lindsay Bond, Mark van Kleunen, Marcel E. Dorken, Marc Stift

## Abstract

Transitions from outcrossing to selfing have been a frequent evolutionary shift in plants and clearly play a role in species divergence. However, many questions remain about the initial mechanistic basis of reproductive isolation during the evolution of selfing. For instance, how important are prezygotic pre-pollination mechanisms (e.g. changes in phenology and pollinator visitation) in maintaining reproductive isolation between newly arisen selfing populations and their outcrossing ancestors? To test whether changes in phenology and pollinator visitation isolate selfing populations of *Arabidopsis lyrata* from outcrossing populations, we conducted a common garden experiment with plants from selfing and outcrossing populations as well as their F1 hybrids. Specifically, we asked whether there was isolation between outcrossing and selfing plants and their F1 hybrids through differences in 1) the timing or intensity of flowering; and/or 2) pollinator visitation. We found that phenology largely overlapped between plants from outcrossing and selfing populations. There were also no differences in pollinator preference related to mating system. Additionally, pollinators preferred to visit flowers on the same plant rather than exploring nearby plants, creating a large opportunity for self-fertilization. Overall, this suggests that prezygotic pre-pollination mechanisms do not strongly reproductively isolate plants from selfing and outcrossing populations of *Arabidopsis lyrata*.

## Background

Mating-system transitions from obligate outcrossing to predominantly selfing have arisen repeatedly across almost all major plant lineages [1]. Up to 15% of seed plants are predominantly selfing and many share a relatively recent common ancestor with outcrossing species [2]. The transition from outcrossing to selfing is usually associated with convergent evolution of a flower morphology that optimizes self-pollination and resource use (e.g. smaller flower size and/or a reduction in pollen, nectar, and scent production), termed the “selfing syndrome” [3–5]. These types of changes in flowering likely contribute to the reproductive isolation of selfing lineages [6], but it is unclear if they or their subsequent effects on pollinators are the main drivers of reproductive isolation in incipient selfing species.

Reproductive barriers are essential to the maintenance of evolutionary independence of parapatric populations (i.e. populations with slightly overlapping ranges). Reproductive isolation can be partial or complete, and most plant species are isolated by a combination of pre- and postzygotic barriers [6–8]. In plants, prezygotic mechanisms are generally more important than postzygotic mechanisms in contributing to the total reproductive isolation of species [6,8–10]. Post-pollination mechanisms such as genetic incompatibilities can cause differences in seed number and/or seed viability, but pre-pollination mechanisms generally contribute more to the total reproductive isolation of plant species [6,9,10]. Although rarely addressed, this principle appears to hold for cases where a shift to self-fertilization has played a part in speciation. For example, in two closely related *Mimulus* species with a contrasting mating system, differences in mean flowering date and floral display contributed the most to reproductive isolation compared to other pre- and post-pollination mechanisms [11]. However, whether the transition to self-fertilization could also promote rapid prezygotic reproductive isolation via changes in floral morphology and associated shifts in pollinator preferences [7] has rarely been studied.

Plant phenological traits and the behaviour of pollinators could readily interact. For example, pollinator behaviour and the number of flowers should both play a large role in determining the opportunity for the flowers of self-compatible plants to be fertilized by a different flower on the same plant (i.e. geitonogamy). Furthermore, geitonogamy could help to reproductively isolate self-compatible individuals. For instance, if pollinators commonly visit multiple flowers on the same individual, it could facilitate higher selfing rates of self-compatible individuals [12]. Different types of pollinators, such as flies vs. bees, could also differ greatly in their pollination strategies [13]. Additionally, if pollinators more commonly visit plants in very close proximity, this could contribute to greater population viscosity and result in more matings among closely related individuals that share the same mating system [14–16]. Furthermore, due to flower attractiveness, pollinators might preferentially and repeatedly visit plants exhibiting a particular mating system type. Alternatively, at the earliest stages of divergence, pollinators might show limited or no ability to differentiate between plants with alternative mating types.

Here, we use *Arabidopsis lyrata* ssp. *lyrata* (L.) to examine the role of differences in phenology and pollinator attraction as mechanisms of reproductive isolation in a recently diverged selfing lineage. In several populations across the range of this normally outcrossing species (multi-locus outcrossing rate: 0.83 < *T* _m_ < 0.99), all plants are self-compatible, have low outcrossing rates, and therefore reproduce primarily through selfing (multi-locus outcrossing rate: 0.09 < *T* _m_ < 0.41) [17–19]. The selfing and outcrossing populations are geographically interspersed, therefore secondary contact following evolutionary divergence in parapatry is likely. Also, the transition to selfing in these populations is thought to have happened < 10,000 years ago because 1) the range now occupied by outcrossing and selfing populations was mostly covered by ice during the last glacial maximum [19], and 2) the selfing populations have not developed a selfing syndrome [20]. This raises the question of whether selfing populations have diverged from their outcrossing ancestors in traits conferring reproductive isolation. Similarly, given that outcrossing and selfing populations are at least partly interfertile and can regularly produce healthy offspring [21,22], F1 hybrids may be a critical factor in determining whether secondary contact would lead to coalescence of the diverged populations or alternatively reinforce their evolved differences.

In a common-garden experiment set within the native range of *A. lyrata*, we simulated two phases of secondary contact between selfing and outcrossing populations. The first phase corresponds to initial contact between parental plants from selfing and outcrossing populations. The second phase corresponds to secondary contact between admixed plants (hybrids between populations) and parental plants. This allowed us to test whether the evolution of selfing has led to pre-pollination isolation through divergence in phenology and/or insect pollinator attraction. Specifically, we asked whether there was reproductive isolation between outcrossing and selfing plants and their F1 hybrids through differences in 1) the timing or intensity of flowering; and 2) pollinator visitation rates and paths. Based on this, we tested whether phenological differences and pollinator behaviour reduced the opportunities for pollen exchange between mating systems. Moreover, as geitonogamy can also contribute to reproductive isolation, we quantified the opportunities for geitonogamy.

## Methods

### Study system

*Arabidopsis lyrata* spp. *lyrata* (L.) is a small, short-lived perennial that is native to North America. It occurs in dry-mesic habitats with shallow soils, such as rock outcrops and sand dunes. Individual plants can produce several stems that terminate in racemes of numerous (>20) small white flowers. The primary pollinators of *A. lyrata* are small solitary bees and hoverflies, which are attracted to the nectar and pollen of the flowers. The ancestral condition in *Arabidopsis lyrata* is self-incompatibility, however the barrier to self-fertilization has broken down in several North American populations [17]. Additionally, many of these newly self-compatible populations have evolved high selfing rates [19]. There is some variation in floral traits, such as flower size and pollen:ovule ratios, among selfing and outcrossing populations, which is primarily explained by population genetic background and not mating system [20].

### Crossing designs

To generate the material needed to simulate secondary contact between diverged selfing and outcrossing populations, we sowed field-collected seeds from 12 North American *A. lyrata* populations with known breeding and mating systems [19] (seeds were kindly provided by Barbara Mable, University of Glasgow). These included six populations characterized as outcrossing (high outcrossing rates, high frequency of self-incompatible individuals, hereafter referred to as SI populations) and six populations characterized as selfing (low outcrossing rates, high frequency of self-compatible individuals, hereafter referred to as SC populations) (Table S1).

In 2012 and 2013, we then produced seeds by manually cross- and self-pollinating up to eight plants per population. To perform the pollinations, we emasculated a flower prior to anther dehiscence, or the same individual in ‘selfed’ crosses, and rubbed a freshly dehisced anther from a haphazardly chosen plant from the same population over its stigma. Progeny were produced with the following cross-types: within SI population (SI-within), and within SC population both by crossing (SC-within) and by selfing (SC-self).

Then to generate the material needed to simulate admixture between the parental populations and their F1 hybrids, we performed a full diallel cross in 2014 and 2015 with six plants of each of the six SI and six SC populations. This cross produced additional progeny of the SI-within and SC-within cross types, as well as the following cross types: between SI population (SIxSI), between SC population (SCxSC), between SC and SI population reciprocally (SIxSC or SCxSI). All crosses were reciprocal and yielded a total of 1032 seed families (Table S2).

### Experimental design of common garden experiment

To test whether differences in phenology and flower-visitor attraction can reproductively isolate plants from selfing populations, we set up a common garden experiment at Trent University in Peterborough, Ontario, Canada. This location is at an intermediate latitude within the geographic range of the source populations (Fig. 1). From March 20 to 22, 2018, for each seed family, up to 50 seeds were sown on a moistened peat-based substrate in one pot. Plants were grown in climate chambers with 11-hour days and a 21°C/18°C day/night cycle at 95% humidity. Between April 18 and May 1, when seedlings had developed at least two true leaves, we transplanted three haphazardly chosen seedlings from each seed family to individual Stuewe and Sons Ray Leach “Cone-tainers” ™ [Tangent, Oregon, USA] with the same peat-based substrate. On May 10, plants were moved to the common garden, prior to any flowering.

**Figure 1:**
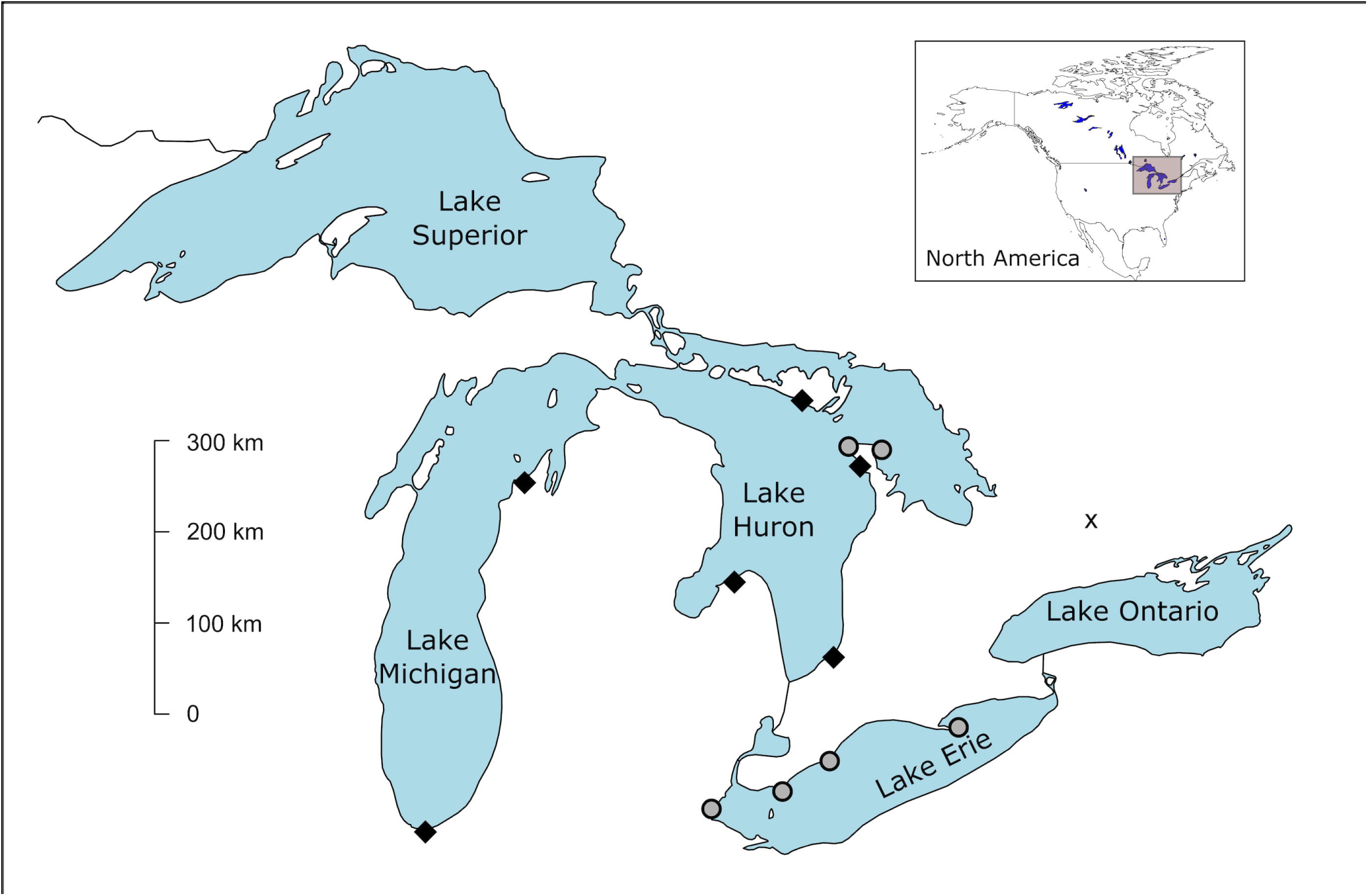
Map of the location of the common garden experiment in relation to the source populations. Gray circles represent selfing populations and black diamonds represent outcrossing populations. The black X represents the location of the common garden at Trent University, Peterborough, Ontario.

Within the common garden, plants were organized in a randomized block design. There were three replicates of three 3×6m blocks. Each of the nine resulting blocks contained between 150-180 individual plants distributed randomly over 180 positions within 9 cone-tainer trays with plants from each seed family and cross type evenly distributed among the blocks. In total, 1509 plants were raised in the common garden. Sample sizes for the cross types were: SI-within (n=172), SC-within (n=175), SC-self (n=65), SIxSI (n=203), SCxSC (n=296), SIxSC (n=314), SCxSI (n=284).

### Phenological data

To test for potential reproductive isolation between the cross types due to differences in phenology, we recorded daily for each plant whether it flowered and how many open flowers it had. Open flowers were defined as flowers with visible reproductive organs (stigma and anthers) and that still had petals attached to the flower. Besides calculating opportunities for pollen-transfer between outcrossing and selfing populations, this allowed us to compare the time to onset of flowering, flowering duration, time of peak flowering, and the maximum flower number (i.e., the number of flowers on the day of peak flowering) for each individual for each of the cross-types.

### Flower visitor observations

To test for differences in insect attraction and flower visitor movements within and between plants, we recorded flowers with GoPro Hero Session^®^ [San Mateo, California, USA] cameras. Specifically, we tested whether there were differences in the potential for geitonogamous selfing (visitor movement within the same plant), and for outcrossing (i.e., visitor movement between plants) within and between mating system. To standardize the recording procedure, 4-6 flowering plants (depending on their size) were taken from their blocks and placed in a tray located at the front of their respective blocks. This method ensured clear video footage of multiple focal plants simultaneously, while minimizing interfering with the visual context of the pollinators provided by the configuration of plants in the block design. To make sure that focal plants had a different set of neighbors for each set of observations, we combined flowering individuals systematically according to their position in the block, going through the block in three different ways: 1) taking consecutive plants in a vertical direction, 2) taking consecutive plants in a horizontal direction, and 3) taking plants from the same position but in different trays. Due to this approach, the cross type of the focal plants combined in the video-frames was random.

We recorded 12-15-minute-long videos that were later trimmed to the central 10 minutes to exclude potential effects of disturbance during starting and stopping the cameras. In total, 500 videos were taken throughout the flowering period, of which a random subset of 140 videos (23.3 hours of video) were analysed by the first author in a random order. In total, these videos included 379 unique individuals (41% of all flowering individuals in the common garden), and 123 plants were observed in multiple videos. For each visitor, we recorded whether it was a solitary bee or hoverfly, the duration of the visit and the path it took (see below). Finer taxonomic identification was not possible due to the video resolution, but we took high quality photographs to identify the most common visitors: hoverflies (Syrphidae) *Eristalis arbustorum, Syritta pipiens, Sphaerophoria* sp. and *Toxomerus marginatus*, and solitary bees from the family Halictidae (kindly identified by Bill Crins, Toronto, Canada).

The path that each visitor took after its initial visit to a flower was recorded to test whether plants from selfing populations received fewer visits than those from outcrossing populations as is expected in selfing plants [23]. Moreover, because pollinators will often focus on exploiting one type of flower and/or floral scent, we tested whether visitors were more likely to move to neighbouring plants with the same mating system than to plants with a different mating system, and whether progeny from crosses between mating system received fewer visits than progeny from crosses within the same mating system. We classified visitor paths as: “away” – the visitor left the video frame after an initial visit; “same” – the visitor visited a second flower on the same individual; or to one of the cross types as defined above (“SI-within”, “SC-within”, “SC-self”, “SIxSI”, “SCxSC”, “SIxSC”, “SCxSI”) – the visitor went to a flower on a different individual. This allowed us to classify the flight paths of the visitors and compare visitation rates among destinations.

### Statistical analyses

All statistical analyses were done in R 3.5.1 [24]. To test if there were differences in the time of peak flowering and duration of flowering between SI and SC cross types (SI-within, SC-within, SC-self) and between within population cross types and between population cross types (SI-within, SC-within, SC-self vs SIxSI, SCxSC, SIxSC, SCxSI), we used Gaussian linear mixed-effects models implemented in lme4 [25] using *cross type* as a fixed effect, and *maternal population* and *paternal population* as random effects. We used “Improper” prior distributions, i.e. distributions with density functions that do not integrate to 1 and are therefore not “proper” probability distributions [26]. Specifically, *p*(β) ∝ 1 was implemented for the model coefficients and *p*(σ^2^) ∝ 1/ σ^2^ for the variance parameters. To obtain the posterior distribution, 5000 values were directly simulated from the joint posterior distribution of the model parameters using the function sim of the R package ‘arm’ [27]. The means of the simulated values from the joint posterior distributions of the model parameters were then used as estimates, and the 2.5% and 97.5% quantiles were used as the lower and upper limits of the 95% credible intervals to make comparisons among cross types.

To test if there were differences in the mean maximum flower number among the cross types, a hurdle model (hurdle function, package ‘pscl’; [28,29]) with a negative binomial distribution that included *maximum flower number* as the response variable and *cross type* as a fixed effect was performed. The hurdle model accounts for the excess number of zero counts in the maximum flower number data. This model specifies one process for zero counts and a separate process for positive counts. The zero counts (flower number as either 0 or 1) were then modelled with a binomial logit model and the positive counts (plants that flowered) with a truncated negative binomial model. The hurdle model also allowed us to calculate the probability that individuals from a cross type would flower.

Pollinator visitation rate (per plant) was analysed separately for the two main visitor classes hoverflies and solitary bees. The cross type ‘SC-self’ was excluded from the analyses of pollinator visitation due to low sample size. To test if there were differences in the frequency of pollinator visits among the cross types, two identical generalized linear mixed-effects models with negative binomial distributions with *number of visits* as the response variable (one model for visits made by hoverflies and another one for solitary bees). The explanatory variables were *cross type* and *flower number* as fixed effects, and *maternal population* and *paternal population* as random effects. In these models, the number of adaptive Gauss-Hermite quadrature points (nAGQ) was set to zero, which optimizes the random effects and the fixed-effects coefficients in the penalized iteratively reweighted least squares step [25]. This results in a faster but less precise parameter estimation for generalized mixed effect models [25]. These models used a log-link function. Improper prior distributions were used, as in the analyses of time of peak flowering and flowering duration.

Pollinator visitation paths were analysed in two ways. The probability that a pollinator would make a certain choice after landing on a flower was analysed with a multinomial logistic regression as implemented in the function multinom in the package ‘nnet’ [30]. *Path* in the multinomial model included all cross types and the same plant (opportunity for geitonogamy) as path options, along with the option of leaving the observation frame. This model included both *cross type* and *flower number* as fixed effects and *path* as the response variable using a logit link function. To further parse the pollinator preference and the effect of flower number and distance between plants in the frame, a conditional logistic regression (function clogit, package ‘survival’; [31]) was performed. The conditional logistic regression was performed separately for hoverflies and solitary bees and included the insect’s selection for any of the cross types in the same video frame as the response variable, as well as *relative flower number, relative distance*, and *cross type* as fixed effects, and finally *switch ID* as the strata. The strata command specifies the group of observations inherent to our video recordings. The strata in this case specifies the group of choice options for each pollinator in each video. Switch ID was defined as: what the insect selected (1) and everything the insect did not select (0) and incorporated information about the distance to the other individuals and the flower number relative to the other individuals. The cross type ‘SI-within’ was used as the baseline as this cross type represents the ancestral condition in *A. lyrata. Relative flower number* and *relative ranked distance* were obtained by dividing by the maximum value within the same video-frame.

### Pollen-transfer probabilities

To examine whether there were differences in the opportunities for self- or outcross pollination between selfing and outcrossing plants, we used the empirical information on phenology and pollinator behaviour to model opportunities for outcrossing between ‘SI-within’ and ‘SC-within’ plants and for geitonogamous pollen movement within plants. Between population cross types were excluded from these analyses. In terms of outcrossing, we were interested in opportunities for pollen exchange within versus between mating types (e.g., whether plants from self-compatible populations of *A. lyrata* had more opportunity to mate with each other than with plants from self-incompatible populations) as a potential mechanism of reproductive isolation. To do this, we used calculations of ***K***_***ij***_ - the “pollen transfer probabilities” outlined in [32]. Here, we use the calculation of ***K*** to refer to the opportunity for mating between plants from different populations in the common garden, based on the overlap in the number of flowers of each mating type per day.

In [32], ***K***_***ij***_ refers to the probability that flowers at the ith position on an inflorescence are pollinated by flowers on other plants at the jth position. Here, we are not interested in the effects of floral position on pollen transfer probabilities, but in the possible effect of mating type. Accordingly, we estimated opportunities for pollen transfer within versus between self-incompatible and self-compatible mating types by calculating the following values of ***K***:

1. ***K***_***ss***_
2. ***K***_***so***_
3. ***K***_***os***_
4. ***K***_***oo***_

where, ***K***_***ss***_ refers to the opportunity for plants from self-compatible populations (SC-within cross type) to fertilize flowers on other plants from self-compatible populations, ***K***_***so***_ refers to the opportunity for plants from self-compatible populations to fertilize flowers on plants from self-incompatible populations (SI-within cross type), and so on.

Brunet and Charlesworth define the probability of pollen transfer between flowers of type ***i*** and ***j*** on day ***c, K***_***ij***_ as: 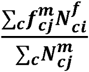 where, 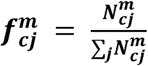.

In the above expressions, the superscripts ***m*** and ***f*** refer to plants in the male and female phases. For *A. lyrata*, which is a simultaneous hermaphrodite, m ***f***, but because we are interested in pollen movement between plants, ***i* ≠ *j***. To calculate mating-type specific values of, for example, ***K***_***ss***_, we calculated ***f***^***m***^ as the proportion of all flowers open per day in the common garden that were from individual plants from self-compatible populations (the SC-within cross type). Therefore, for this calculation, the value of the numerator, 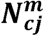, was calculated only for those plants. Plants from self-incompatible populations were included in the calculation of ***N***^***m***^ for ***K***_***os***_ and ***K***_***oo***_. For all values of ***K***, all plants were included in the calculation of the denominator of 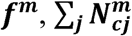.

The calculated values of ***K*** are frequency dependent - a small group of plants of one mating type surrounded by plants of the opposite mating type would have more opportunities for between, rather than within mating-type pollen transfer. Because we were specifically interested in opportunities for pollen transfer driven by phenology, not frequency, we used bootstrapping to generate 200 randomly sampled, equal-sized populations of plants of each mating type for the calculation of ***K***. For each mating type, we sampled 200 plants (with replacement) of each cross type for inclusion in each calculation of ***K***.

Two values of ***K*** refer to within mating-type fertilization opportunities and two of them to between mating-type fertilization opportunities. To evaluate whether plants from populations of the two different mating types (self-compatible versus self-incompatible) differed in the proportion of within-versus between mating-type pollen transfer opportunities, we used linear, mixed models for each set of bootstrapped values of ***K***. Population was included as random effect in these models. In the absence of phenological differences between plants from self-compatible and self-incompatible populations, the expected value of the parameter estimate for the mating-type effect is 0. Accordingly, to test whether plants representing the two mating types differed in their opportunities for within-versus between mating-type pollen transfer, we tested whether the distribution of parameter estimates from each set of bootstrapped values of ***K*** differed from 0 using a two-tailed *t*-test.

The opportunity for geitonogamous self-pollination is determined by the number of simultaneously open flowers per plant and the likelihood that pollinators will move from one flower to another on the same plant. Videos of pollinator movements provide per-population estimates of that likelihood. We calculated the opportunity for geitonogamous pollen transfer, ***G***_***c***_, for plants with ***n*** open flowers as a geometric series of the likelihood of within-plant pollinator movement ***x***. That is, for each day ***c, G***_***c***_ = ***x*** + ***x***^**2**^ + ***x***^**3**^ + … + ***x***^***n*−1**^. The total opportunity for geitonogamous pollen transfer over the flowering season was calculated as **Σ**_***c***_***G***_***c***_.

## Results

### Phenology

Of the 1509 plants in the common garden, 938 flowered (62%). The main flowering period lasted six weeks from June 1 to July 14, although a few individuals flowered later (nine individuals flowered a second time and 10 individuals flowered for the first time as late as September) (Fig. S1). The probability of an individual flowering varied by cross type. SI-within and SC-within cross types did not strongly differ from each other in the probability of flowering (55% and 61%, respectively; CrI overlapping; Fig. 2a), however the probability of flowering of the SC-self cross type (29%) was substantially lower (Fig. 2a). So, while progeny formed by selfing flowered less, merely having the ability to self did not substantially decrease the probability of flowering when compared to individuals from outcrossing populations. The F1 hybrid cross types did not differ from the within-population cross types in the probability of flowering (54%-64%; CrIs overlapping; Fig. 2a), with the exception of the SIxSI cross type being more likely (83%) to flower than the other F1 or within-population cross types (Fig. 2a). Additionally, the direction of the cross for SIxSC and SCxSI hybrid F1 crosses did not have an obvious effect on flowering probability, as both cross types had similar probabilities (63% and 64% respectively) for flowering (Fig. 2a).

**Figure 2:**
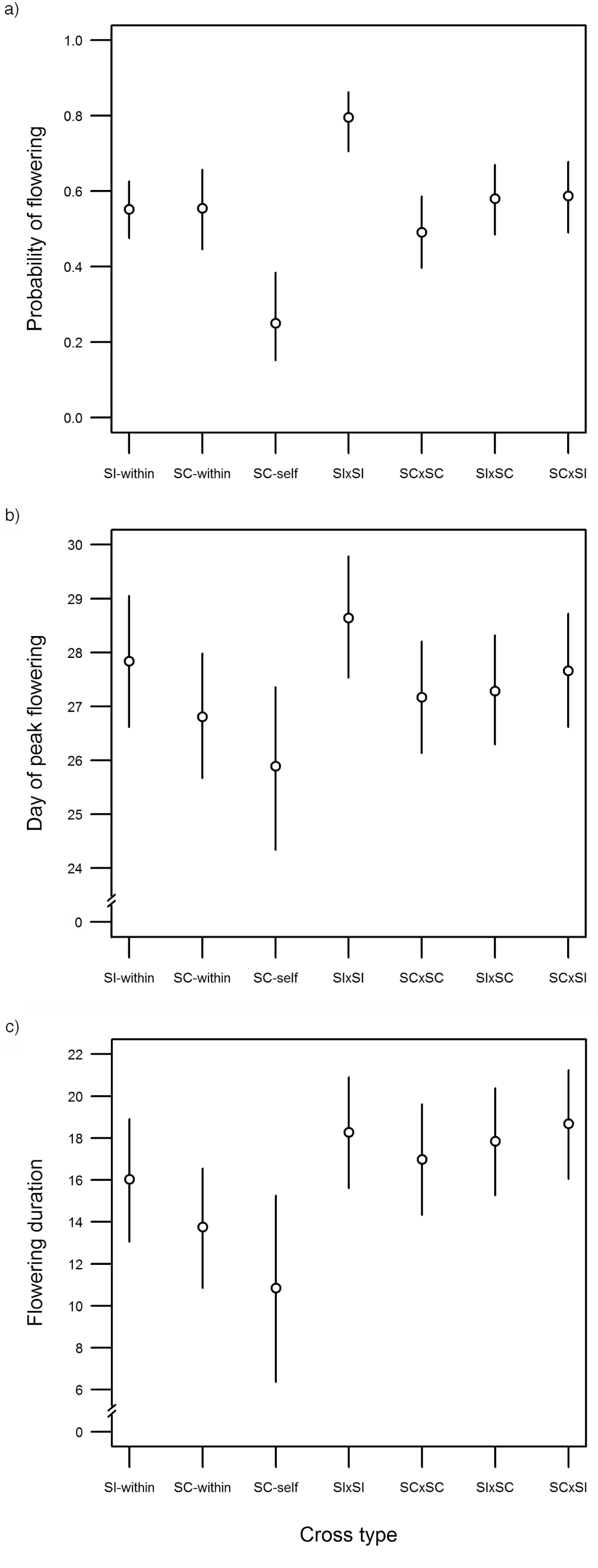
Panel of effect plots for the differences in phenological traits among the cross types (n=1509). a) The probability of flowering per cross type. Fitted values are obtained from the hurdle model. b) The day of peak flowering per cross type. This was calculated as the day where each individual had the highest number of flowers during the total flowering period. Fitted values obtained are from the Gaussian linear model that included *day of peak flowering* as the response variable. c) The flowering duration (days) per cross type. Fitted values obtained are from the Gaussian linear model that included *flowering duration* as the response variable. Vertical bars in all figures represent the 95% credible intervals. SI-within= within outcrossing population cross type, SC-within= within selfing population cross type, SC-self= selfed, SI= self-incompatible, SC= self-compatible. The probability of flowering and time of peak flowering varied among the cross types. Flowering duration did not strongly differ among the cross types.

The time of peak flowering differed among the cross types. While there were no strong differences in the time of peak flowering between the SI-within and SC-within cross types, the SC-self cross type peaked in flowering about one to two days earlier (average = 27.6 days) than the other within or F1 cross types (however, CrIs overlapped with all cross types except SIxSI; Fig. 2b). The day of peak flowering did not differ between SIxSC and SCxSI plants, indicating that cross direction did not influence the time of peak flowering (CrIs overlapping; Fig. 2b). There were also no strong differences in peak flowering between the within-population and F1 cross types (CrIs overlapping), with the exception that the SIxSI cross type tended to reach peak flowering one to two days later (Fig. 2b). Similarly, flowering duration did not strongly differ among the cross types (Fig. 2c), but the SC-self cross type tended to have a shorter duration (6-15 days) than the other within population cross types (11-19 days) or F1 cross types (15-21 days) (Fig. 2c).

For flowering plants of all cross types, the mean maximum number of open flowers on a single day ranged from 10.8 to 13.6 flowers. After correcting for zero-inflation in the maximum flower number model, there were no strong differences in maximum flower number among the cross types.

### Pollinator visitation

There were no differences in flower visitation between the within population cross types or between the within population cross types and the F1 hybrids (all CrIs overlapping; Fig. 3a, Fig. 3b). Solitary bees and hoverflies were the predominant visitors, and they had similar visitation frequencies and no clear pattern of preference for any of the cross types (compare Fig. 3a and 3b). The behaviour of both types of pollinators appeared to facilitate geitonogamous self-pollination, as ∼50% of the movements between flowers were to a different flower on the same plant (Fig. 4). When cases of a pollinator visiting another flower on the same plant were not considered, the odds of an initial visitor moving to a plant in the frame decreased by 89% (solitary bees) and 94% (hoverflies) for each unit increasing relative distance (significantly negative odds-ratios for relative distance; Table 1). In other words, pollinators were more prone to visit the nearest plant, regardless of the cross type or the number of flowers on the neighbouring plant.

**Table 1.** Summary of results from the conditional logistic regression model that analyzed whether *relative flower number, relative distance*, or *cross type* influenced the pollination path. Cross type ‘SI- within’ was used as the baseline. Hoverflies: n=546, number of events = 142, Likelihood ratio test= 62.16 on 7 df, p=<.001*. Solitary Bees: n=541, number of events=140, Likelihood ratio test= 45.24 on 7 df, p=<.001*. Symbols and abbreviations used in the column headings: SE= standard error; z= Wald statistic. For both hoverflies and solitary bees, relative plant distance had the greatest influence on pollinator choice. There was no strong preference for any of the cross types or individuals with more flowers. Significant effects are highlighted in **bold**.

| Visitor | Fixed effect | Odds Ratio | SE | z | Pr(>\|z\|) | 95% confidence interval |  |
| --- | --- | --- | --- | --- | --- | --- | --- |
| **Solitary bees** | Relative Flower Number | 1.10 | 0.32 | 0.29 | 0.77 | 0.59 | 2.04 |
|  | Relative Distance | **0.11** | **0.38** | **-5.73** | **<.001\*** | **0.05** | **0.24** |
|  | SC-within | 2.01 | 0.39 | 1.80 | 0.07 | 0.94 | 4.30 |
|  | SIxSI | 1.32 | 0.39 | 0.71 | 0.48 | 0.62 | 2.82 |
|  | SCxSC | 0.96 | 0.39 | -0.12 | 0.91 | 0.44 | 2.07 |
|  | SIxSC | 0.80 | 0.36 | -0.64 | 0.52 | 0.40 | 1.60 |
|  | SCxSI | 0.91 | 0.38 | -0.25 | 0.81 | 0.43 | 1.91 |
| **Hoverflies** | Relative Flower Number | 0.90 | 0.34 | -0.32 | 0.75 | 0.46 | 1.75 |
|  | Relative Distance | **0.06** | **0.43** | **-6.56** | **<.001\*** | **0.03** | **0.14** |
|  | SC-within | 1.06 | 0.45 | 0.14 | 0.89 | 0.44 | 2.55 |
|  | SIxSI | 1.09 | 0.41 | 0.21 | 0.83 | 0.49 | 2.44 |
|  | SCxSC | 1.70 | 0.40 | 1.34 | 0.18 | 0.78 | 3.71 |
|  | SIxSC | 1.03 | 0.40 | 0.07 | 0.95 | 0.47 | 2.27 |
|  | SCxSI | 0.83 | 0.41 | -0.44 | 0.66 | 0.37 | 1.86 |

\* indicate p-values <.05. The 95% confidence interval is the confidence interval of the odds ratio. *Relative flower number* and *relative distance* were estimated relative to the other individuals in the observation. These variables were transformed to range between 0 and 1.

**Figure 3:**
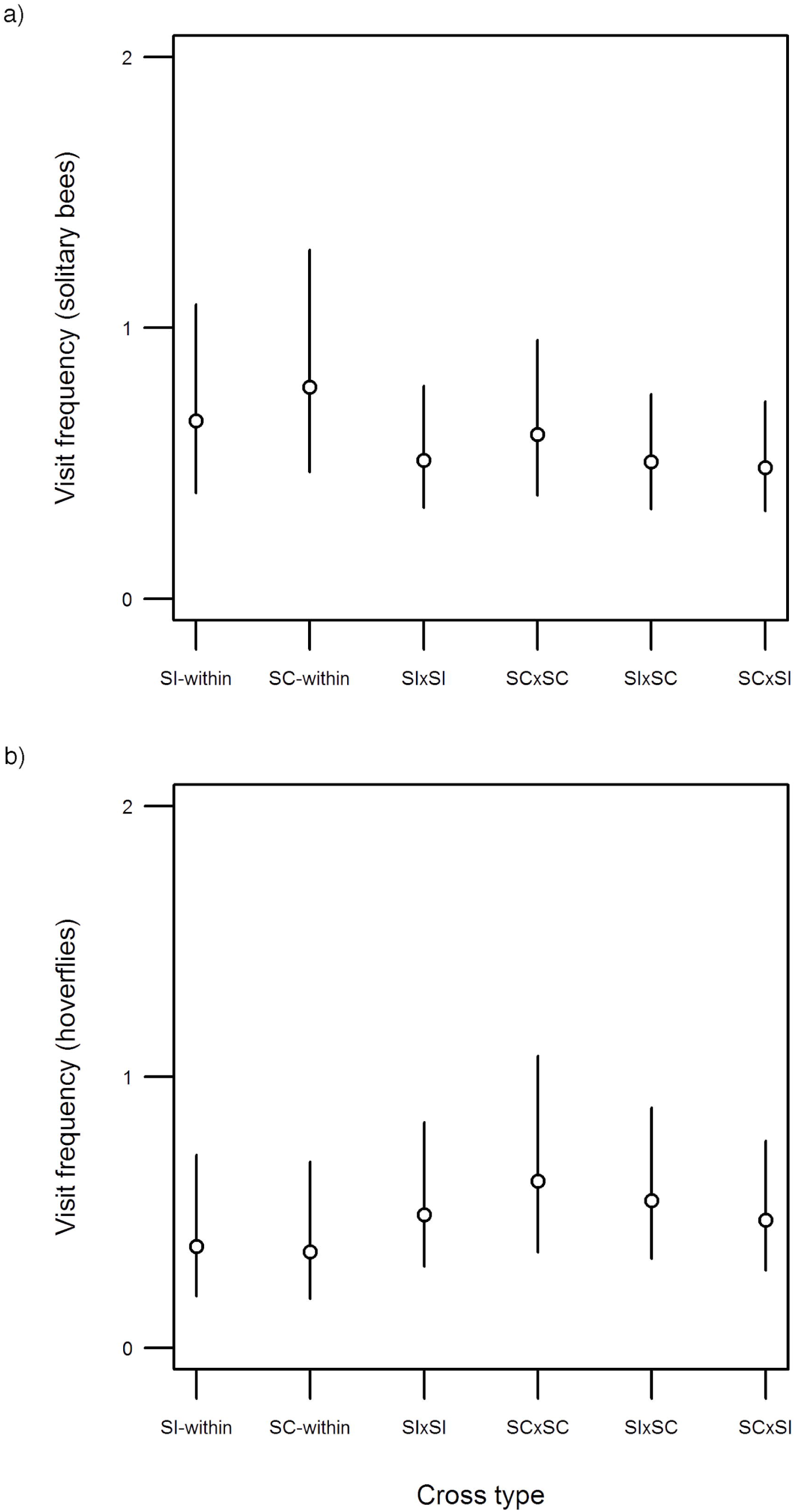
Differences in pollinator visitation (number of visits) by a) solitary bees and b) hoverflies among the cross types (n=502). The fitted values obtained are from the generalized linear models with negative binomial distributions described in the methods. Vertical bars represent the 95% credible intervals. SI-within= within outcrossing population cross type, SC-within= within selfing population cross type, SI= self-incompatible, SC= self-compatible. The number of visits did not strongly differ among the cross types and pollinator identity had a minor influence on the number of visits. Solitary bees made slightly more visits overall.

**Figure 4:**
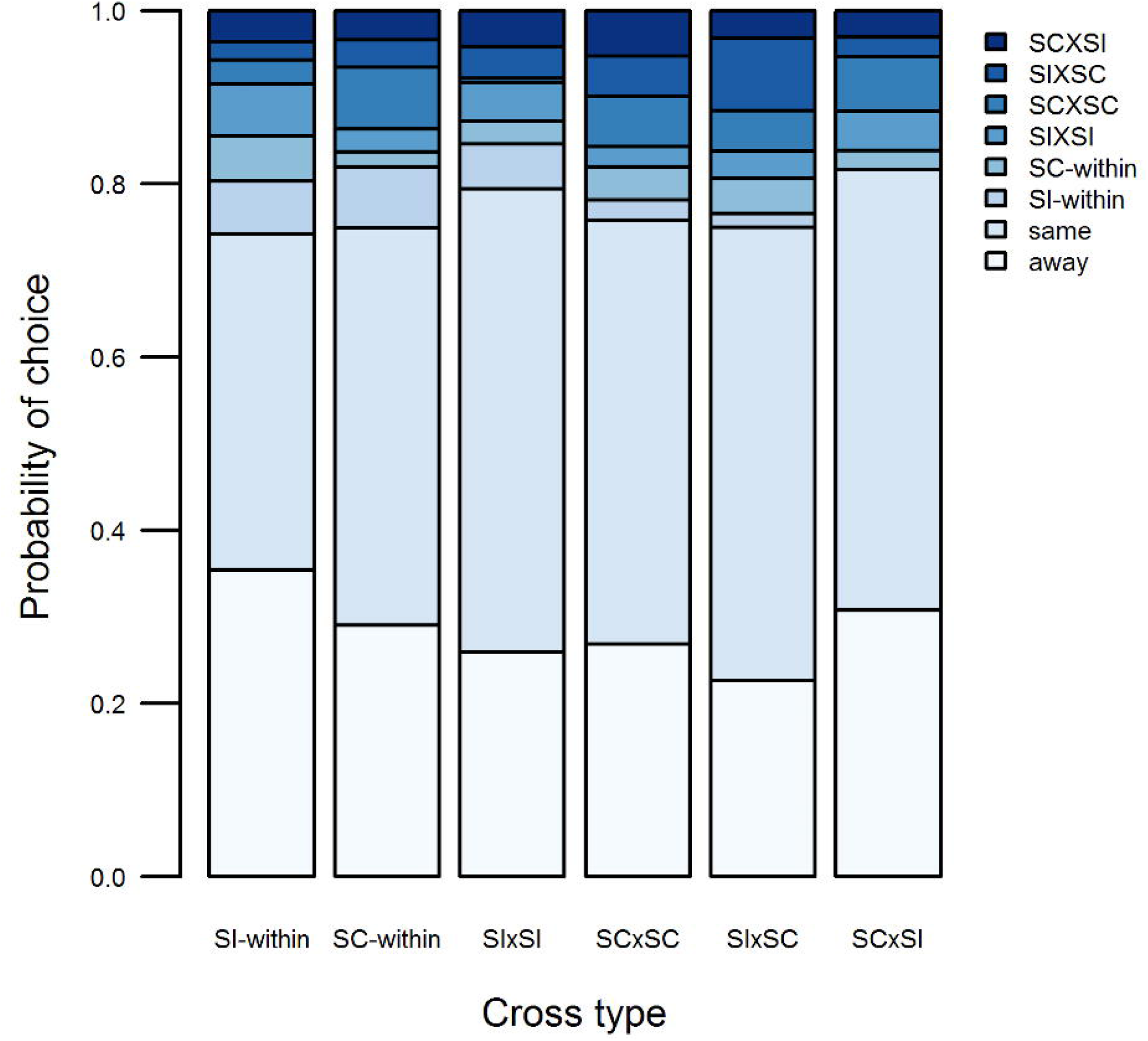
Stacked bar plot representing the probability of pollinators making a choice to visit an individual of any cross type after visiting an individual of a certain cross type. Probabilities were obtained from the multinomial model (and thus corrected for the number of available flowers on each plant in the array). Pollinators could also choose to visit a flower on the same plant (“same”) or to leave the experimental setup (“away”). The cross types on the x-axis represent the cross type of the initial visit. The stacked blue bars represent the probability of that cross type being selected after a visit to the cross type on the x-axis. SI-within= within outcrossing population cross type, SC-within= within selfing population cross type, SI= self-incompatible, SC= self-compatible.

### Opportunities for pollen-transfer

Based on a bootstrapping approach that integrated the timing and intensity of flowering throughout the entire flowering period, the opportunity for between vs. within cross type pollen-transfer was nearly equal both for the SI-within and SC-within cross type (Fig. 5). In other words, slight shifts in phenology and flowering intensity (Fig. S1) are unlikely to lead to reproductive isolation. Additionally, a similar approach that also took into account the pollinator paths showed that the opportunity for geitonogamy arising from plants having multiple flowers open simultaneously and the high frequency of within plant movement of pollinators (Fig. 4) did not differ between SI-within and SC-within plants.

**Figure 5:**
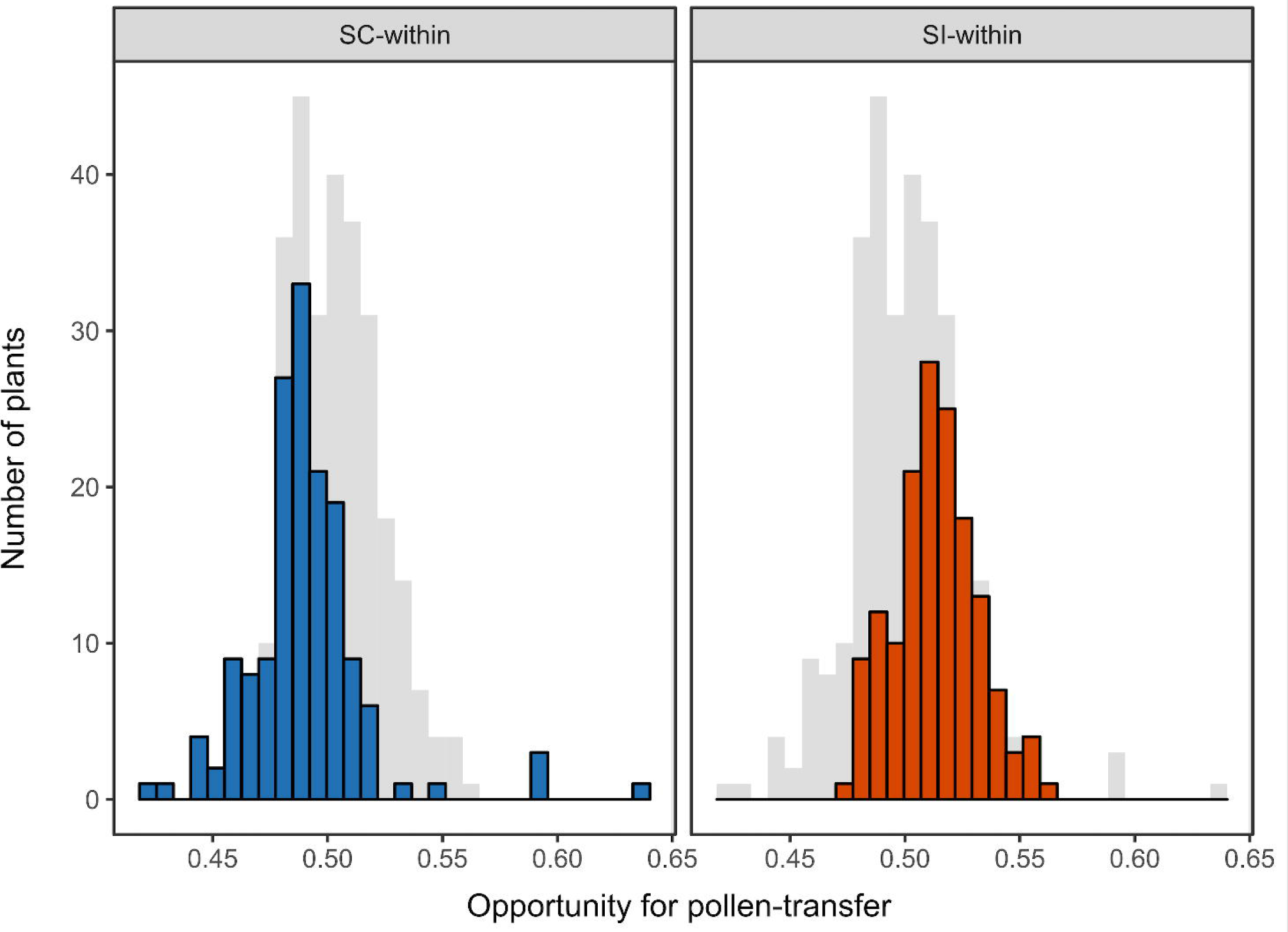
Representative bootstrapped run of the distribution of the self- and outcross pollen transfer opportunities for self-compatible and self-incompatible individuals. The parameter estimate for the difference in transfer-probability for the shown run was 0.024, and thus close to the mean value of 0.020 across all the bootstrapped samples. There were no significant differences in the opportunities for pollen transfer.

## Discussion

Our common garden experiment simulating secondary contact between outcrossing and selfing populations showed that phenology largely overlapped between plants from outcrossing and selfing populations. There were also no differences in pollinator preference related to mating system. Regardless of mating system, pollinators tended to move between flowers on the same plant, thus facilitating opportunities for geitonogamy. Our models of pollen-transfer probabilities, which integrated differences in phenology (timing and intensity of flowering), revealed equal opportunities for pollen-transfer within- and among mating systems. Together, this suggests that prezygotic pre-pollination mechanisms do not strongly reproductively isolate plants from selfing and outcrossing populations of *Arabidopsis lyrata*. However, because plants simultaneously open multiple flowers, and pollinators predominantly move from flower to flower on the same plant, our data suggest that there is a large opportunity for geitonogamy in this system. Such geitonogamy could isolate selfers to some extent.

### Pollinator visitation

Our results show that differences in pollinator preference do not play a large role in the reproductive isolation of the newly diverged selfing populations. We found no differences in the preferences of the two main pollinator types, hoverflies and solitary bees, among the cross types. Additionally, when given a choice pollinators preferred to stay on the same individual versus exploring nearby plants regardless of cross type. When they did choose to visit another plant, pollinators preferred to go to the closest plant regardless of how many flowers it had or what cross type it was. This lack of pollinator preference could be due to the pollinators simply being generalists, as previous studies have shown that hoverflies, for instance, are not very choosy with respect to the plants they visit [13]. Additionally, there could be a lack of floral differences because selfing individuals still need pollinators to transfer their pollen as they are not completely autonomous [20]. Overall, pollinator preference seems to be playing little role in differentiating the selfing from outcrossing populations of *A. lyrata*.

Nevertheless, the behaviour of the pollinators could favour selfing for several other reasons. For instance, we found that pollinators often visit different flowers on the same individual, irrespective of mating system. This should provide ample opportunity for within-individual pollen transfer [3,33], and thus for self-compatible individuals to self-fertilize through geitonogamy. Moreover, when pollinators moved between plants, they mainly moved between nearby individuals. Given that *A. lyrata* seeds have no mechanisms to promote dispersal, and plants can produce over 1000 seeds per season, neighbouring plants could be highly related to each other [34]. As a consequence, the observed behaviour of pollinators could cause mate limitation in self-incompatible plants, making the transfer of cross-pollen rare and/or mainly from incompatible partners (e.g., from relatives that share S-alleles). In self-compatible plants, on the other hand, geitonogamy may help overcome this mate limitation and provide reproductive assurance [35]. Even without mate limitation, where geitonogamy does not provide reproductive assurance but rather results in seed discounting, selfing could still be favoured due to the associated inherent transmission advantage [36]. The transmission advantage would favour selfers if inbreeding depression is low. Indeed, the relatively low estimates of inbreeding depression for our study populations [37] imply that such transmission advantage alone could be sufficient to drive the evolution of selfing. There is usually a low frequency of self-compatible individuals in outcrossing populations [19,38]. Since the observed pollinator behaviour should promote geitonogamous selfing, it remains enigmatic why selfing has not evolved in all North American populations of *A. lyrata*.

### Potential consequences of admixture

We found that F1 hybrid cross types had a similar phenology and pollinator visitation as the parental cross types. Earlier studies have shown that hybrids between outcrossing and selfing plants can be intermediate for phenological traits. For instance, in the genus *Clarkia*, hybridization between selfing and outcrossing populations resulted in floral characteristics and flowering times that were intermediate between the parental populations [39]. Our results show similar relationships between F1 hybrids and the parental populations in regard to flowering probability and time of peak flowering. This suggests that in a scenario of secondary contact, F1 hybrids would likely function as a bridge to further gene exchange between selfing and outcrossing plants, which could potentially lead to the parental populations merging [40]. Whether the resulting admixed populations will maintain a mixed mating system [2], or evolve to become predominantly selfing or outcrossing remains to be tested. Initially, as inbreeding depression tends to be low [37,41], selfing may be favoured due to the associated inherent transmission advantage. However, on longer timescales, expression of drift load may select against selfing as shown in selfing populations of *A. lyrata* [22](but see [42]). It would therefore be of interest to monitor the performance and mating system of admixed populations over multiple years.

### Conclusions

Our common garden experiment showed that, although pollinator behaviour may isolate selfers by promoting geitonogamy, outcrossing and selfing *A. lyrata* populations are only weakly reproductively isolated via pre-pollination mechanisms. These findings differ from findings in other systems with a recent transition to selfing (e.g., [43,44]). The weak isolation between selfing and outcrossing populations of *A. lyrata* is likely because its transition to selfing is even more recent, and has not led to evolution of a selfing syndrome [20]. Future studies could investigate if reproductive isolation due to prezygotic pre-pollination mechanisms are larger in natural populations, giving specific attention to parapatric selfing and outcrossing populations. Moreover, to what extent other mechanisms such as niche differentiation and genetic incompatibilities contribute to reproductive isolation remains to be investigated.

## Acknowledgements

Christina Steinecke and Ryan Holt helped with data collection. Colleen White helped with transplanting. Bill Crins taxonomically identified pollinators. Fränzi Korner-Nievergelt and Pius Korner-Nievergelt (oikostat GmbH) helped with parts of the statistical analyses. Barbara Mable provided the source material (seeds) from wild populations. Yan Li, Ekaterina Mamonova, and Sina Konitzer-Glöckner helped with the crosses to produce the seeds used in the experiment.

## Data Accessibility

Data will be made available from the Dryad Digital Repository upon acceptance.

