## Supplemental Figure 1 for "Limited phenological and pollinator-mediated isolation among selfing and outcrossing *Arabidopsis lyrata* populations"

Fig S1: The number of open flowers (per population) per day over the total flowering period. Blue lines represent selfing populations and orange lines represent outcrossing populations. Only populations for which we had more than 10 plants that flowered are plotted.

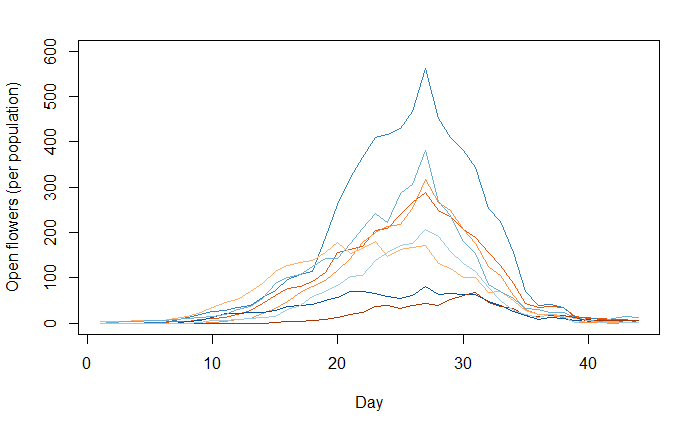
