## Supplemental Table 1 for "Limited phenological and pollinator-mediated isolation among selfing and outcrossing *Arabidopsis lyrata* populations"

Table S1. Background information for populations used in this study. The populations are a subset of those used in Foxe et al (2010). Outcrossing rates (T_m_) and frequencies of self-incompatible (SI), self-compatible (SC) and partially compatible (PC) individuals are taken from Foxe et al. (2010).

| **Population** | **Location, State/Province, Country** | **Population Coordinates** | | **T_m_** | **SI** | **SC** | **PC** |
| --- | --- | --- | --- | --- | --- | --- | --- |
| **IND** | Indiana Dunes National Lakeshore, Indiana, USA | N 41˚37'17" | W 87˚12'44" | 0.99 | 0.86 | 0.14 | 0.00 |
| **KTT** | Kitty Todd State Nature Preserve, Ohio, USA | N 41˚37'14" | W 83˚47'15" | 0.31 | 0.00 | 1.00 | 0.00 |
| **LPT** | Long Point Provincial Park, Ontario, Canada | N 42˚34'47" | W 80˚23'15" | 0.13 | 0.00 | 1.00 | 0.00 |
| **MAN** | Manitoulin Island, Ontario, Canada | N 47˚39'54" | W 82˚15'52" | 0.83 | 0.88 | 0.13 | 0.00 |
| **PCR** | Port Crescent State Park, Michigan, USA | N 44˚00'15" | W 83˚04'26" | 0.98 | 0.75 | 0.00 | 0.25 |
| **PIN** | Pinery Provincial Park, Ontario, Canada | N 43˚16'08" | W 81˚49'53" | 0.84 | 1.00 | 0.00 | 0.00 |
| **PTP** | Point Pelee National Park, Ontario, Canada | N 41˚55'40" | W 82˚30'51" | 0.09 | 0.00 | 1.00 | 0.00 |
| **RON** | Rondeau Provincial Park, Ontario, Canada | N 42˚15'41" | W 81˚50'47" | 0.28 | 0.00 | 1.00 | 0.00 |
| **SBD** | Sleeping Bear Dunes National Lakeshore, Michigan, USA | N 44˚56'20" | W 85˚52'13" | 0.94 | 0.75 | 0.00 | 0.25 |
| **TC** | Tobermory Cliffs, Bruce Peninsula National Park, Ontario, Canada | N 45˚14'30" | W 81˚31'03" | 0.18 | 0.13 | 0.88 | 0.00 |
| **TSS** | Tobermory Singing Sands, BPNP, Ontario, Canada | N 45˚11'33" | W 81˚35'02" | 0.91 | 1.00 | 0.00 | 0.00 |
| **TSSA** | Tobermory Singing Sands Alvar, BPNP, Ontario, Canada | N 45˚11'27" | W 81˚35'26" | 0.41 | 0.38 | 0.50 | 0.13 |
