## Supplemental Table 2 for "Limited phenological and pollinator-mediated isolation among selfing and outcrossing *Arabidopsis lyrata* populations"

Table S2. The crossing design that generated the seeds used for the experiment. Population is abbreviated as ‘pop.’ and mating system is abbreviated as ‘MS’.

| **Cross Type** | **Crossing Formula** | **Seed Families** | **Cross Type Produced** |
| --- | --- | --- | --- |
| Within pop. | 12 pop. x 3 cross-combinations x 2 MS x 2 directions | 144 | SI-within, SC-within |
| Within pop. and selfed | 12 pop. | 96 | SI-within, SC-within, SC-self |
| Between pop. | 6 sets x 15 pop. pairs x 2 MS x 2 directions | 360 | SIxSI, SCxSC |
| Between MS | 6 sets x 36 SI-SC pop. pairs x 2 directions | 432 | SIxSC, SCxSI |
